# Integration of memory and sensory information in skilled sequence production

**DOI:** 10.1101/2025.09.10.675426

**Authors:** Amin Nazerzadeh, Medha Porwal, J. Andrew Pruszynski, Jörn Diedrichsen

## Abstract

Sequential movements rely on two information sources: external sensory cues and internal memory representations. Although often both sources jointly drive sequential behavior, previous research has primarily examined them in isolation. To address this, we trained participants to perform sequences of rapid finger presses in response to numerical cues. Sensory influence was measured by varying the number of visible cues, and memory influence by comparing repeating and random sequences. Early in learning, participants integrated sensory and memory information: repeating sequences were performed more quickly when more cues were visible. After learning, when repeating sequences were predictable with certainty, participants relied solely on memory and ignored sensory cues. However, when this certainty was manipulated by introducing occasional violations within repeating sequences, participants reverted to integrating memory with sensory cues. We propose a computational model that successfully predicted both speed and accuracy of individual presses. Critically, this model relied on the assumption that multiple movements are planned independently of each other. This independence assumption was then validated by examining response patterns to isolated violations in repeating sequences. Finally, we provide evidence into how sequence memories can be flexibly deactivated and reactivated in response to these violations. Together, these results reveal how brain dynamically integrates sensory and memory information to produce sequences of movements.

## Introduction

Human movements rarely occur in isolation. In many daily activities, we perform long and complex sequences of tightly interlocking movements. These sequences can be guided by external cues in the environment, or by internal memory representations formed through practice (Fitts, 1964; Adams, 1987). Consider a pianist playing a musical piece. They may play the melody from the sheet music, or they may ignore the visual information and rely solely on memory, having internalized the piece through prior practice. Often, however, both sources of information are available. In such cases, the pianist may integrate the visual information from the sheet music with their internal memory, allowing for a smoother and more accurate performance.

How does such integration happen during sequence production? Previous literature often characterized sequence production as either purely sensory-driven or as purely memory-driven (Welford, 1968; Karni *et al*., 1995; Willingham, 1998; Wu, Kansaku and Hallett, 2004; Simó, Krisky and Sweeney, 2005), with performance switching from one mode to the other with practice (Adams, 1987; Proctor and Dutta, 1995; Picard, Matsuzaka and Strick, 2013; Wymbs and Grafton, 2015; Mizes *et al*., 2023). The fact that sequence learning is gradual, however, suggests that the learners spend a significant portion of training in intermediate stages where both sensory input and memory guide production. Moreover, even in highly practiced sequences, performance is flexible and can take into account new sensory information (Hikosaka *et al*., 1999; Xu *et al*., 2022). Thus, the integration of memory and sensory information during sequential movements is likely a very important aspect of movement skill, which remains poorly understood.

One notable exception, in which the combined use of external sensory cues and memory has been extensively explored, is probabilistic sequence learning (Lewicki, Hill and Bizot, 1988; Howard and Howard, 1997; Schvaneveldt and Gomez, 1998; Jiménez and Méndez, 1999). However, this line of work has mainly been conducted using the Sequential Reaction Time Task (SRTT), where participants only receive visual information about one response at a time. Therefore, learning in this task likely reflects mostly the cognitive anticipation of the next elements in a sequence (Keele *et al*., 1995; Willingham *et al*., 2000; Wong *et al*., 2015), rather than the formation of a motor sequence memory. For learning the latter, participants must have knowledge of multiple future elements of a sequence at the same time, allowing them to link their movements into a coordinated sequence (Krakauer *et al*., 2019; Kashefi, Diedrichsen and Pruszynski, 2025).

To study the integration of sensory information with sequence memory, we used a Discrete Sequence Production (DSP) task where participants performed sequences of finger presses in response to numerical cues (Verwey, 1996, 2001). A key manipulation was the *visible horizon*, which determined how many upcoming presses participants could see. This allowed us to assess how future visual information was used for action planning (Ariani *et al*., 2021). Moreover, some sequences were repeated such that participants could form an internal memory representation. In contrast, no memory was available for random sequences. By examining performance for repeating sequences across different visible horizons, we aimed to characterize how sensory input and memory are integrated during sequence production and how this integration evolves with practice.

We showed that once repeating sequences were fully learned, participants relied entirely on memory to guide their performance. With a small injection of uncertainty, however, participants reverted to integrating sensory information with their memory of the sequence. We proposed a simple computational model that reproduced the participants behavior. This model was based on the strong simplifying assumption that participants plan multiple future movements independently. We tested this assumption explicitly by examining how participants responded to isolated violations within learned sequences. Finally, we provided some first insights into how the memory of a specific sequence is activated and deactivated during performance.

## Results

Participants were instructed to perform sequences of finger presses on a keyboard-like device by pressing the corresponding keys in response to numerical cues presented on a monitor (**Fig. 1A**; see methods). In Experiment 1, participants practiced two unique repeating sequences alongside random sequences over the course of three days. At the beginning of each trial, a symbol indicated the upcoming sequence type (**Fig. 1A**). For repeating sequences, the participants therefore could rely on both memory and visual cues, whereas for random sequences, they needed to rely on numerical cues alone. We further varied the number of visible digits at each time point (Visible Horizon, **Fig. 1B**). In Horizon 1, only the visual cue of the next to be pressed (current) digit was available. For longer Horizons (2, 3 or 7) cues for future presses were available. Additionally, for repeating sequences, we introduced masked trials in which all digits remained masked, forcing participants to rely on memory only.

**Figure 1.**
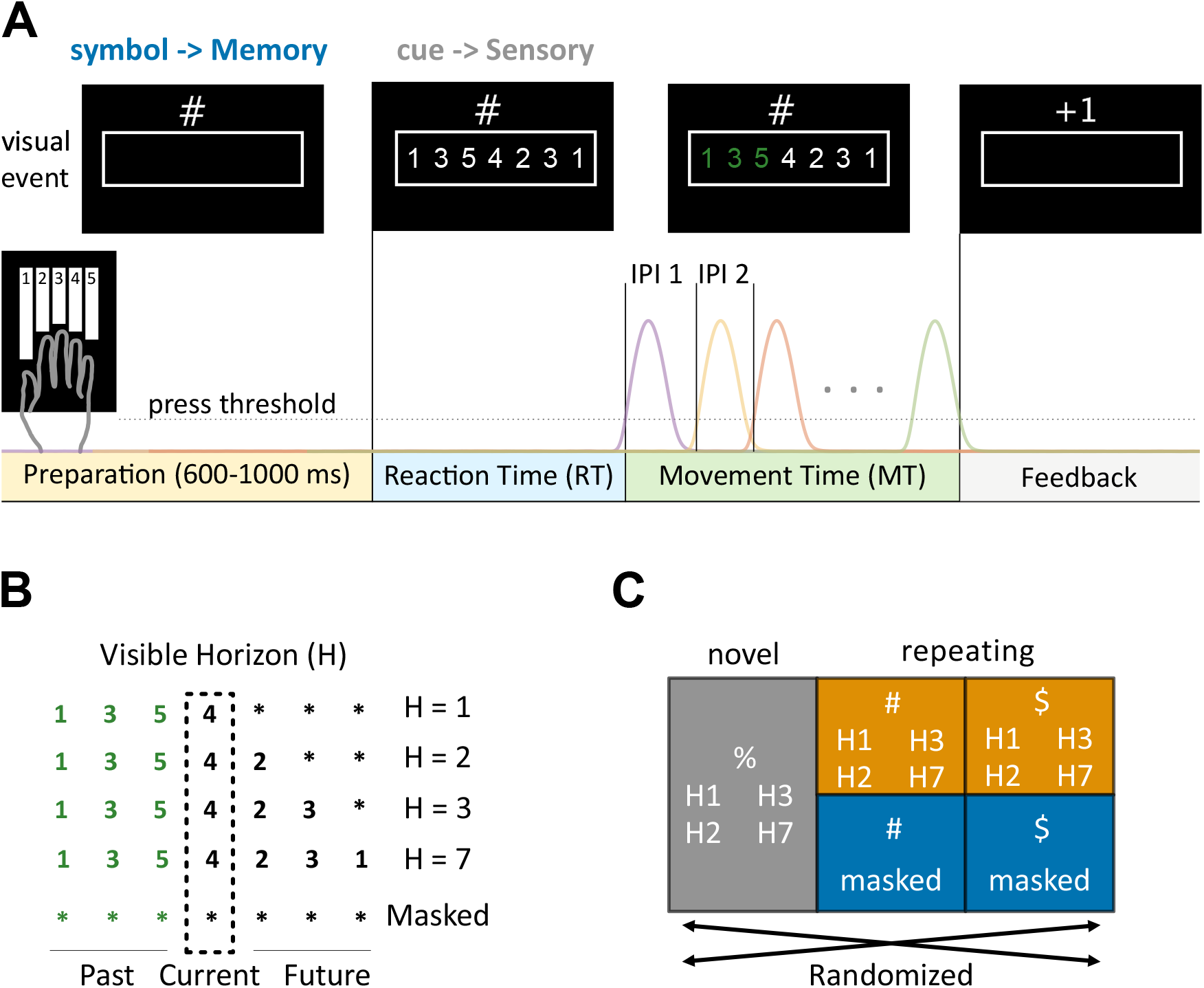
Experiment 1 methods. A) Timeline of a single trial. During the preparation phase a memory symbol (#, $, %) was presented. It was followed by numerical cues that indicated the keys to be pressed. Movement time started when the force of the first press (violet trace) exceeded the press threshold and ended when the last press (green trace) was completed. B) Visible horizon conditions. An example screen just before the current digit (dashed box) was pressed. Asterisks indicate masked digits. The horizon conditions (H1-H7) differed in the number of future cues available at each moment. When the current digit was pressed, the next digit was unmasked. In the masked condition, all digits remained masked. C) Experimental Design: Each block was counterbalanced with random (%), sequence 1 (#), and sequence 2 ($).H1, H2, H3, H7 were counterbalanced within sequence types. In repeating sequences, masked and unmasked were counterbalanced. Trials were randomized within a block.

### For random sequences, participants plan 2-3 items ahead

To understand how participants used the available sensory information without any memory, we compared movement times across different visible horizons for random sequences. As expected, having a longer horizon reduced movement time. Averaged across days, the difference between H1 and H2 (*t*_13_ = 18.952, *p* = 7.482e-11), as well as the difference between H2 and H3 (*t*_13_ = 4.572, *p* = 5.240e-4) was significant (**Fig. 2B**). This suggests that participants used the information of up to 3 digits into the future to plan random sequences (Ariani *et al*., 2021).

**Figure 2.**
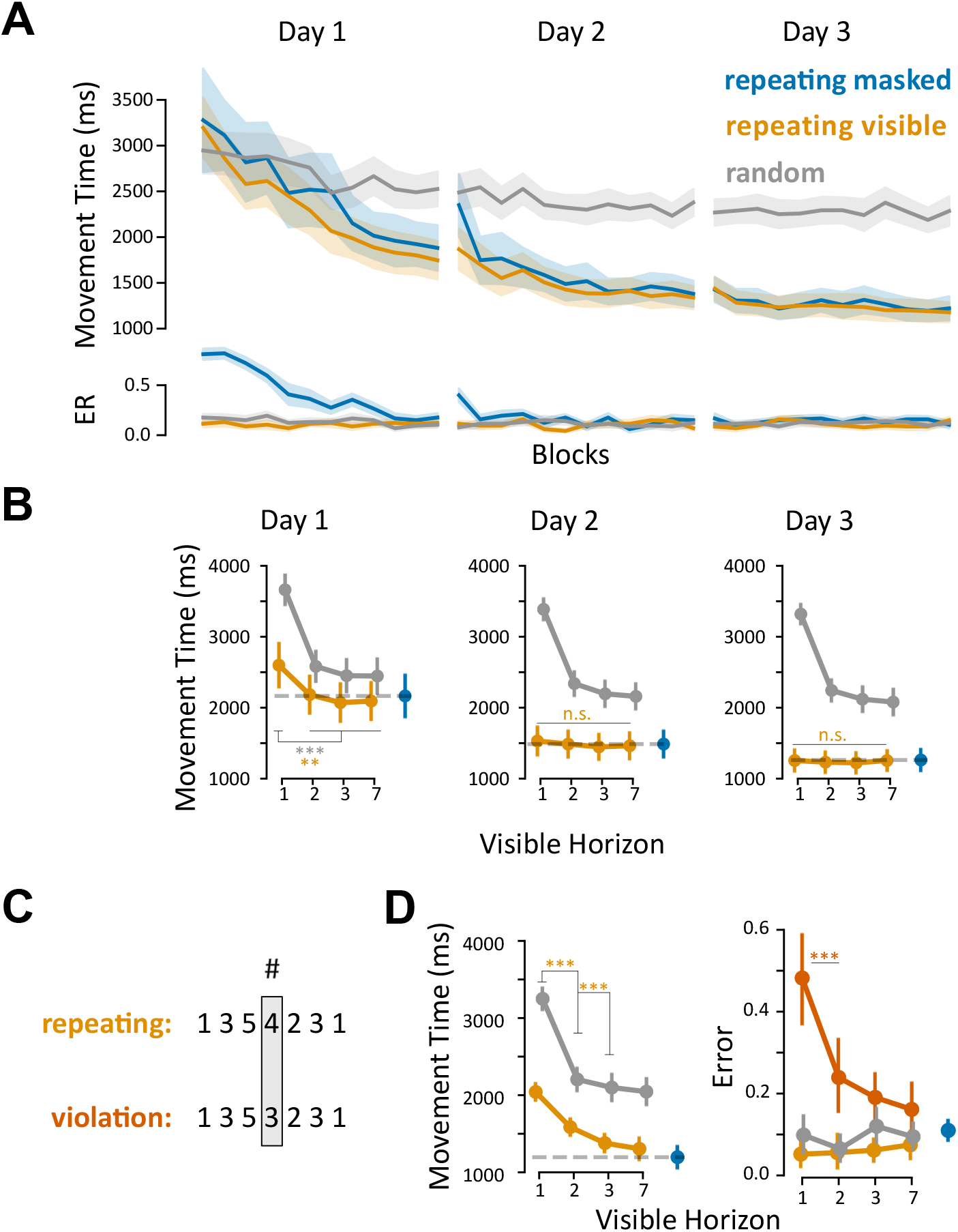
Experiment 1. A) Group-averaged movement times (top) and error rates (bottom) across three days of learning for random (gray, averaged across H1-H7), repeating visible (yellow, averaged across H1-H7), and repeating masked (blue) sequences. B) Group-averaged movement times as a function of visible horizon (x-axis) across three days of learning. The repeating masked condition (blue) is shown on the right for comparison. C) Example violation trial on day 4: in these trials, a single cue from the repeating sequence was changed (shaded box). The same sequence symbol (#, shown on top) was displayed in both conditions. D) Group-averaged movement times (left) and error rates (right) for each visible horizon on day 4. Error rates for violation condition are shown in red. For all plots: error bars show the standard error of the mean across participants. ** and *** show p < 0.01 and p < 0.001.

### After initial learning, repeating sequences are performed from memory only

On day 1, participants were presented with their individually selected repeating sequences for the first time. Consequently, the error rate in the masked condition – where no visual cues were available – was initially very high and only reduced over the course of the day (comparing the first and the last three blocks, *t*_13_ = 12.159, *p* = 1.783e-8; **Fig. 2A**). When visual cues were visible, random and repeating sequences started out with similar error rates and movement times. Over the course of the day, however, repeating sequences were executed more quickly than random sequences (*t*_*13*_ = 6.130, *p* = 3.600e-5). Breaking up the performance of repeating sequences by horizon (**Fig. 2B**), we found faster performance for longer horizons (*F*_3,39_ = 16.482, *p* = 4.500e-7), consistent with the results in random sequences. These findings demonstrate that participants integrated memory and the available sensory information early in learning.

In contrast, after training on day 3, participants performed the repeating sequences from memory only 20 ± 12 ms slower than when visual cues were available, a non-significant difference (*t*_13_ = 1.598, *p* = 0.134; **Fig. 2A**). Additionally, the effect of the horizon became non-significant (*F*_3,39_ = 1.176, *p* = 0.331). This suggests that late in learning, participants relied entirely on memory and ignored the available sensory information.

### Reappearing of memory and sensory integration under uncertainty

In contrast to many sequences in everyday life, repeating sequences on day 3 could be fully predicted from the sequence symbol. To investigate how the participant’s behavior would change under more uncertain conditions, we introduced occasional violation trials on the fourth day. In these trials, the numerical cues differed from the sequence indicated by the sequence symbol at one random position (**Fig. 2C**; see methods). With violations occurring in 3 out of 8 trials, the upcoming cues were only partially predictable based on the sequence symbol.

In response to this increased uncertainty, participants changed their behavior on the non-violation trials (**Fig. 2D**, orange). In contrast to day 3, movement times on day 4 showed a significant benefit for longer horizons (*F*_3,39_ = 72.144, *p* = 5.694e-16), indicating a return to horizon-dependent performance. At the same time, participants still performed repeating sequences faster than random ones (averaged across horizons, *t*_*13*_ = 10.928, *p* = 6.367e-8), suggesting continued use of memory. These findings suggest a strategic shift from purely memory-driven performance to integration of memory and sensory under uncertainty.

This integration provided a behavioral advantage: While error rates on violation trials were much higher than on non-violation trials (*t*_13_ = 8.815, *p* = 7.611e-7), that error rate decreased with longer horizons (*F*_3,39_ = 16.066, *p* = 5.903e-7; **Fig. 2D**). This confirmed that participants used future sensory cues to resolve the conflict between memory and sensory input.

### A model of memory and sensory integration for single responses

These findings raise a key question: what kind of computation enabled participants to resolve conflicts between memory and sensory input when future cues were available – while still preserving their speed advantages in longer horizons? One might imagine that with a longer horizon, participants can detect the conflict earlier. At the same time, longer horizons lead to faster performance, leaving less time to resolve the conflict and increase the risk of errors.

To understand how these factors may combine to result in the observed behavior, we first built a model that captured the behavior in the simplest condition, Horizon 1, where only the visual cue for the current press was available. We then extended this model to make predictions in conditions with Horizon > 1, where multiple future cues were available.

In the Horizon 1 condition (Experiment 1; **Fig. 3C, right column**), reaction times (RTs) were faster when memory and sensory input were compatible (repeating) compared to when no memory was available (*t*_13_ = 8.027, *p* = 2.154e-6). In contrast, when memory and sensory input were in conflict (violation), RTs increased relative to random trials (*t*_13_ = 7.968, *p* = 2.335e-6) and accuracy dropped (violation vs others: *t*_13_ = 8.421, *p* = 1.270e-6). These observations suggest that at the single-response level, memory and sensory input reinforce each other when aligned, and compete when in conflict.

**Figure 3.**
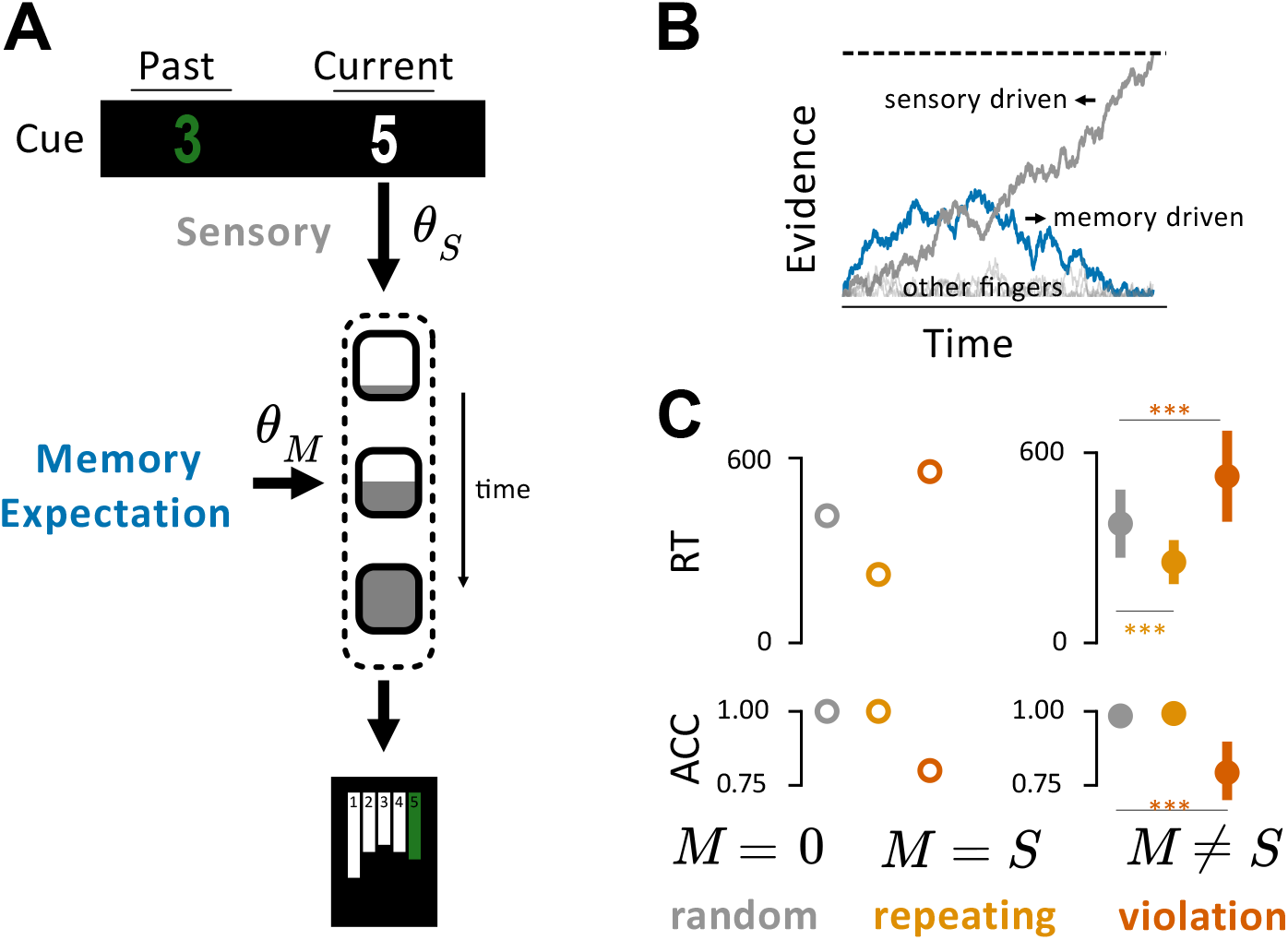
Decision model of a single response. **A)** Sensory information from the cue jointly with the memory expectation drive an evidence accumulation with different weights (*θ*_*S*_ *and θ*_*M*_). This process of evidence accumulation is illustrated by the box gradually filling over time. When the decision process reaches a threshold, a press is issued. **B**) Example evidence accumulation for a press on which memory expectation and sensory input are in conflict. Evidence state of the finger driven by memory (blue) and of the finger driven by sensory (grey) are shown as a function of time. Evidence states of other fingers only fluctuate with noise. The decision threshold is indicated by the dashed line. c) Reaction times (RT) and accuracy (ACC) of presses at positions where cue changed in the violation trials. Hollow circles indicate model’s predictions (left), and solid circles indicate group-averaged data for the Horizon 1 condition, Experiment 1, Day 4 (right). Error bars show the standard error across participants. ** and *** show *p* < 0.01 and *p* < 0.001.

To capture these observations, we modeled a single response as an evidence accumulation to a decision bound (Usher and McClelland, 2001; Bogacz *et al*., 2006). Each finger (i.e., response option) accumulated evidence driven by both memory and sensory inputs (see methods; **Fig. 3A**). The first candidate that reached threshold was chosen as the response. The additive effect of the two inputs drove the process to threshold faster, explaining the RT benefit when memory and sensory are compatible (**Fig. 3C**). To explain the slowdown in RT in violation trials, we introduced lateral inhibitions between evidence states of finger (matrix A, see methods). In a violation trial (**Fig. 3B**), the finger indicated by memory initially gained evidence faster, but was ultimately overtaken by the finger indicated by the sensory cue. The final model (see methods) captured correctly the faster performance on repeating trials, as well as the slow-down and increased error rate on violation trials.

### A model of memory and sensory integration for sequences

In the next step we extended the model to the sequential context when more than one cue was available. We previously observed that random sequences were performed more quickly when the visible horizon was larger. This suggested that participants were planning for future presses at the same time they were planning for the current press.

To account for this, we extended the single press decision model by assuming that the evidence accumulation for up to three presses into the future can occur in **parallel** and **independently** (**Fig. 4A**, see methods). Starting with the parameters fitted to Horizon 1 data, we introduced new input weights for future presses (**Fig. 4A**). These input weights were estimated using the non-violation trials from the Horizon > 1 conditions for both random and repeating sequences. The model correctly fitted the faster MTs for larger horizons (**Fig. 4B**), because evidence for future presses can accumulate while the current press is being selected. By the time the current press reaches threshold, the next two presses are partially selected, resulting in an overall reduction in production time.

**Figure 4.**
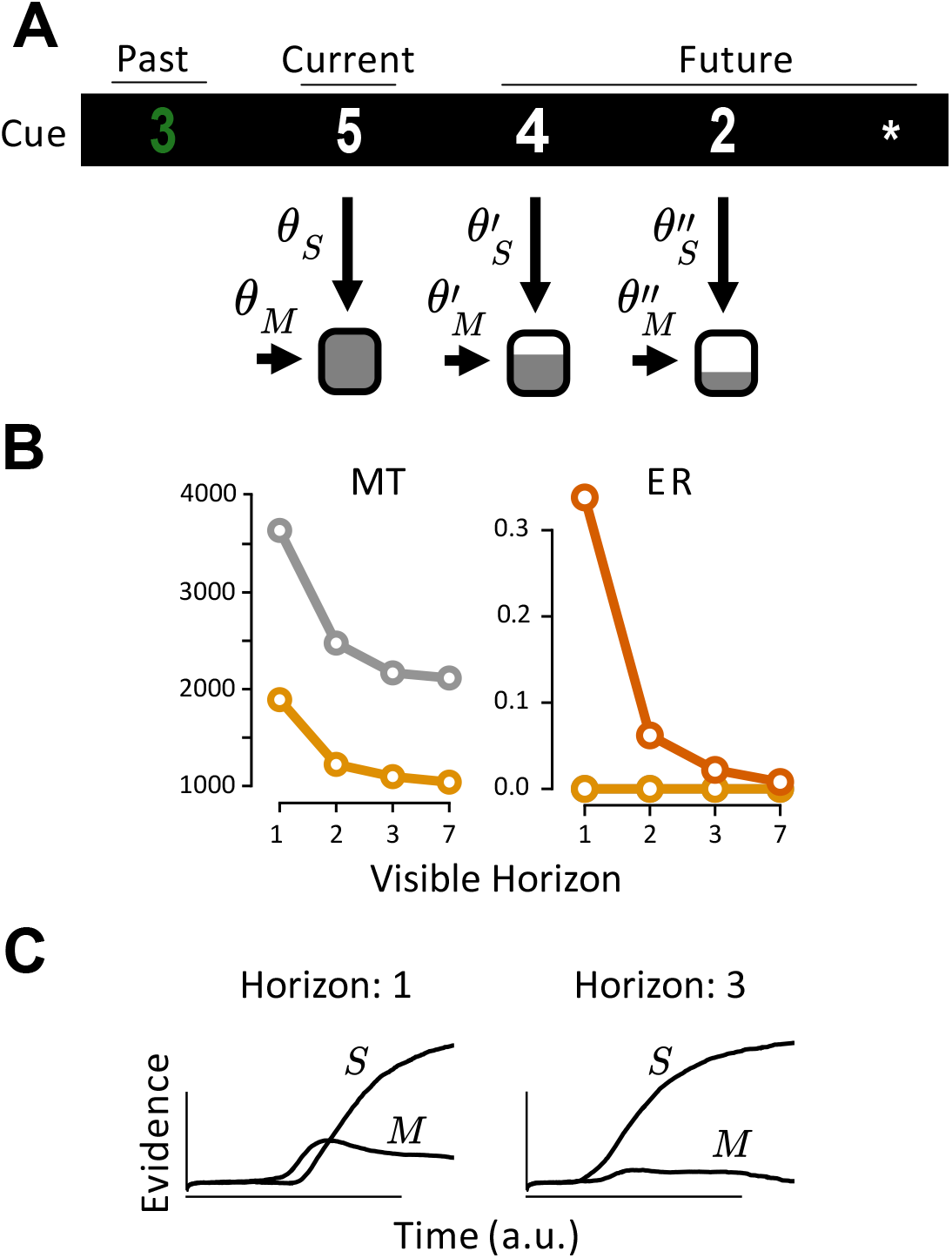
The decision model in sequential contexts. A) An example trial in H3 showing parallel and independent sensory and memory integration for the current and two future presses. Memory and sensory inputs drive these decision processes with different weights. B) Fitted decision model’s production movement times (left) and error rates (right) across visible horizons. C) Evolution of evidence state on a press where S and M are conflicting for H1 (left) and H3 (right). Other fingers are not shown for brevity. Time axis is aligned to the start of the trial.

We then tested whether this model could accurately predict the behavior in violation trials for Horizon > 1. Without refitting the model to these data, it predicted a decrease in error rates for violation trials at longer horizons, consistent with the behavioral results (data: **Fig. 2D**, model: **Fig. 4B**). This benefit arose from the model’s ability to resolve conflicts by looking ahead: in Horizon 1, the sensory information was available only after the memory-driven process had started accumulating evidence for the incorrect finger, increasing the chance that this finger reached the threshold first (**Fig. 4C**, left). In contrast, in Horizon 3-7, sensory cues were available earlier, allowing evidence for the correct finger to accumulate due to the sensory-driven process, which in turn inhibited the incorrect memory-driven finger. (**Fig. 4C**, right).

### Independent planning predicts no slowdown before violation press

Together, our modeling results demonstrated that a parallel evidence accumulation framework can explain faster performance when both memory and future sensory information (Horizon > 1) are available and captures how participants resolve memory-sensory conflicts in the sequential setting.

We then tested one important simplifying assumption that we made when constructing the model: that the evidence accumulation processes for the current and the future presses occurred **independently**. The independence assumption predicts that a conflict between sensory and memory input in a future press leaves the decision processes for earlier presses unaffected. As a result, the model predicts that while inter-press intervals should be slower on the press where violation happens, they should not differ between violation and non-violation trials on preceding presses (**Fig. 5A** Model). Indeed, any slowdown prior to the violation would indicate interdependence between decision processes.

**Figure 5.**
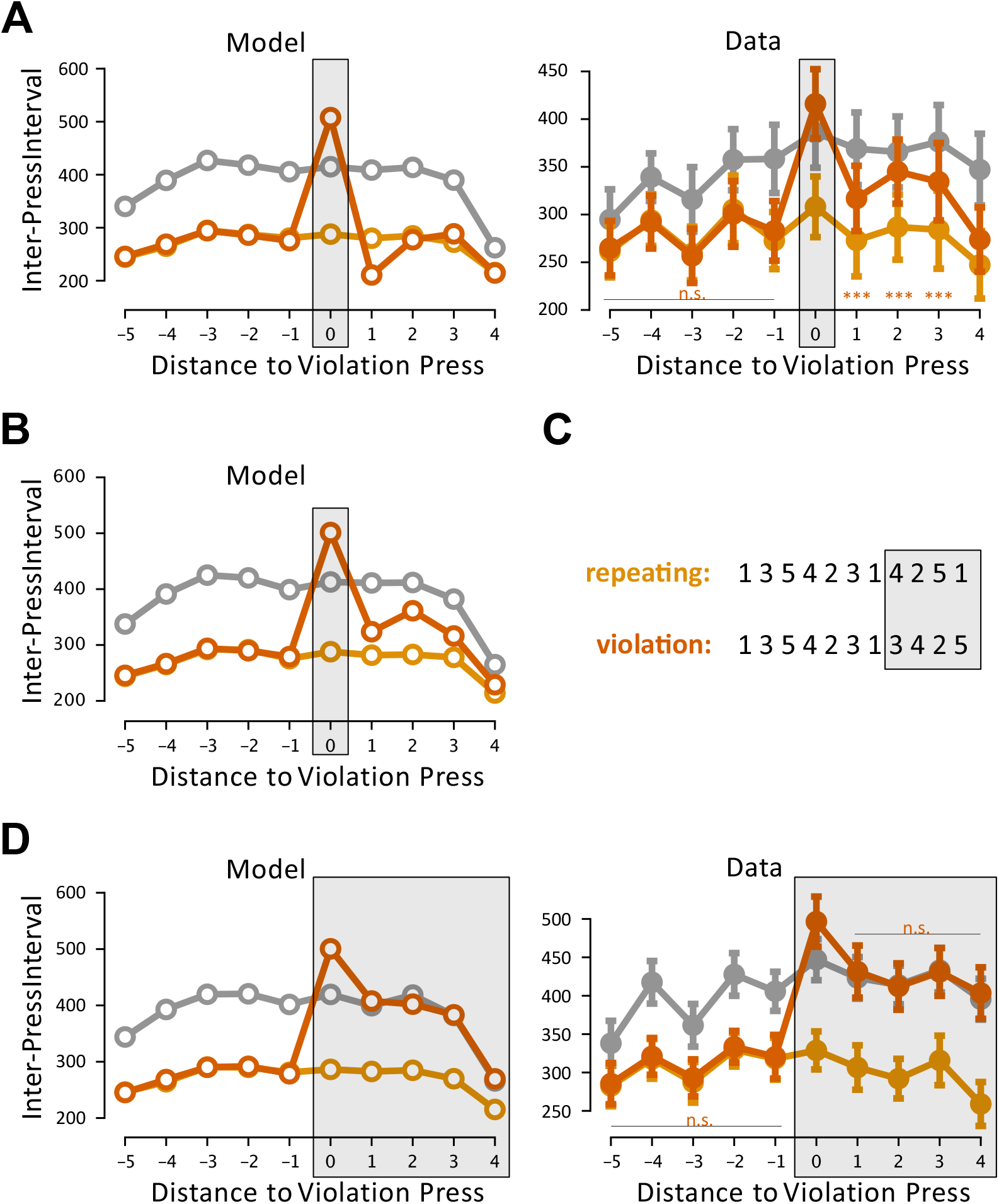
Experiments 2 and 3. A) Inter-press intervals aligned based on their distance to the positions where change happens in the violation trials in Experiment 2 (shaded boxes). Negative distances show preceding presses and positive distances show following presses. Model predictions are shown with hollow circles (left) and Experiment 2 data are shown with solid circles (right). B) Model predictions with fast memory activation and slow deactivation in Experiment 2. C) Example violation trial in Experiment 3. All cues after a certain position (shaded box) are in conflict with previously practiced repeating sequence cues. D) Inter-press intervals aligned to where change happens in violation trials in Experiment 3. Model predictions are shown with hollow circles (left) and Experiment 3 data are shown with solid circles (right). For all plots: Error bars show the standard error of the mean across participants. *** shows *p* < 0.001.

Because Experiment 1 did not have a sufficient number of violation trials to test this prediction, we conducted Experiment 2 (see methods), training participants on longer sequences (11 digits) with visible horizons 2, 3, 4, and 11. As established before, participants used sensory future cue information in repeating sequences (**Supplementary Fig. 1**). Critically, we examined the press at which a violation occurred, as well as the three preceding presses. Averaged across horizon conditions, we again observed a substantial slowdown on the press where violation happened (108 ± 28 ms, *t*_13_ = 3.919, *p* = 2.398e-3). Importantly, the IPIs for preceding presses did not significantly differ between violation and non-violation trials (*3 before:* −3 ± 1 ms, *t*_13_ = −1.864, *p* = 0.089; *2 before*: −5 ± 3 ms, *t*_13_ = −1.769, *p* = 0.105; *1 before*: 10 ± 5 ms, *t*_13_ = 2.013, *p* = 0.069, **Fig. 5A** Data). These results support the model’s prediction that conflicts in future presses do not disrupt earlier presses and suggest that sequential finger presses can be planned independently, despite temporal overlap in their planning.

### Deactivation and re-activation of sequence memory

Although our model correctly captured participant behavior before and during the violated press, it failed to predict the behavior after the violation. Specifically, the model predicted that performance should return to the level of the non-violation trials immediately after the conflict (**Fig. 5A** Model). Contrary to this prediction, participants showed persistent slowdown on the four presses following a violation, compared to non-violation trials (*1 after*: 44 ± 9 ms, *t*_13_ = 4.732, *p* = 6.178e-4; *2 after*: 58 ± 13 ms, *t*_13_ = 4.459, *p* = 9.637e-4; *3 after*: 50 ± 11 ms, *t*_13_ = 4.777, *p* = 5.740e-4; *4 after*: 26 ± 13 ms, *t*_13_ = 2.056, *p* = 6.428e-2; **Fig. 5A** Data). This occurred despite the cues following the violation being fully compatible with the expected memory of the sequence, suggesting that the violation had a longer-lasting effect on performance.

One explanation is that participants temporarily reduce their reliance on memory following a conflict. In this view, the violation acts as a signal to downregulate memory-based planning, leading to a performance more similar to a random sequence. To model this process, we introduced a mechanism in which the memory input weights (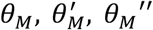) are dynamically updated on each press. Specifically, when a conflict occurs on the violation press, memory input weights are decreased (deactivated). When the sensory information becomes consistent again with the memory on following presses, memory input weights are increased (reactivated). This dynamic was realized in the model by two time-constants determining the rates of memory activation and deactivation. Fitting these to participant data (see methods), resulted in a fast deactivation time constant (0.88 ± 0.06) and a slow reactivation time constant (0.33 ± 0.1). This indicated that memory disengages rapidly in response to conflicts but takes longer to recover. This updated model successfully captured the persistent slow-down observed after the violation press (**Fig. 5B**).

Alternatively, the observed post-violation slowdown could reflect a surprise-induced global performance decrement (Botvinick *et al*., 1999; Vassena, Deraeve and Alexander, 2020), rather than memory-specific disengagement. According to this post-surprise slowdown explanation, violations should elicit a general cautious control response that transiently slows down any motor performance (for example by raising the decision threshold). Therefore, if post-violation cues were altered to follow a random pattern, performance on these presses should be slower than on random trials, where no surprise occurs. In contrast, the memory deactivation model predicts when post-violation cues follow a random pattern, behavior should match that of a random trial because memory no longer contributes after the violation (**Fig. 5D** Model).

To test the two predictions, we conducted Experiment 3 (see methods), which was similar to Experiment 2 but differed in that in violation trials, all cues after a certain position were changed from the repeating sequence – i.e., after a violation, the sequence never returned back to the originally learned sequence (**Fig. 5C**). On the first violation press, because of the conflict, participants were 57 ± 26 ms slower than random presses (*t*_11_ = 2.146, *p* = 0.055). However, on the subsequent presses, the performance in violation trials was not statistically different from the random sequences (*1 after*: 5 ± 12 ms, *t*_11_ = 0.418, *p* = 0.684; *2 after*: 12 ± 18 ms, *t*_11_ = 0.694, *p* = 0.502; *3 after*: −10 ±19 ms, *t*_11_ = −0.524, *p* = 0.611; *4 after*: 6 ±10 ms, *t*_11_ = 0.653, *p* = 0.527; **Fig. 6D** Data). These observations favor a memory deactivation rather than a non-specific control response as the explanation for the slower performance following a sequence violation.

## Discussion

Everyday sequences of movements are usually neither fully predictable nor completely random. Therefore, sequential behaviors in everyday life require integrating internal memories and external sources of information. We studied this ability in the context of rapid sequences of finger presses. By systematically manipulating availability of memory and visual cues, we found that early in learning, participants integrated both sources of information to guide performance. Once sequences were well learned and could be predicted with certainty, participants ignored sensory cues and relied solely on memory. These findings highlight a learning-dependent shift from an integrative mode to a memory-driven mode of sequence production (Fitts and Posner, 1979; Logan, 1988).

However, not all well practiced sequences in life are executed purely from memory. Flexibility in movement generation is essential, allowing the brain to adapt sequences of movements based on dynamic environmental or sensory input (Rosenbaum, 2009). While different neural mechanisms and circuitries have been proposed to underlie this flexibility (Hikosaka *et al*., 1999; Xu *et al*., 2022; Mizes *et al*., 2023), the influence of varying amounts of sensory input on this flexibility has not been fully explored.

In our task, we occasionally violated the expected cues in well-learned sequences with varying amounts of sensory input, introducing uncertainty that increased production errors and required flexible adjustments in performance. By integrating sensory information in planning multiple presses concurrently, participants leveraged the availability of future cues in longer horizons to reduce errors and achieve flexibility.

### Decision model captures main characteristics of sequential sensory and memory integration

We were able to explain our findings using a sequential decision model. Decision models have been successful in explaining how sensory inputs are integrated with motor memories to control individual movements (Trommershäuser, Maloney and Landy, 2008; Berniker and Kording, 2011; Wolpert and Landy, 2012). Here, we extended this type of model to understand how this integration occurs when multiple actions are performed in rapid succession. In our model, we propose that decision processes for multiple presses occur in parallel. This simple assumption reproduced the observed increase in production speed when visual information about upcoming items were available. Interestingly, just by this simple assumption, the model predicted how participants integrated the visual cue of future presses to resolve the conflict in violation trials and minimized their errors.

### Independent planning of sequential movements

A surprising insight from our model was that participants’ behavior could be explained even when assuming independent decision processes for each future movement. We tested this assumption by introducing isolated violations into learned sequences. The resulting violation affected only the performance on the violated press, but not the preceding presses – supporting the idea of independent planning.

The notion of independent planning contrasts with numerous behavioral findings suggesting interdependencies in sequence planning. Behavioral patterns such as chunking (Rosenbaum, Kenny and Derr, 1983; Abrahamse *et al*., 2013), co-articulation in reaching (Ramkumar *et al*., 2016; Kashefi *et al*., 2024; Kashefi, Diedrichsen and Pruszynski, 2025), path tracking (Bashford *et al*., 2022), or speech production (Kent and Minifie, 1977) point to coordinated planning across elements of a sequence.

In contrast there is electrophysiological evidence that movement plans for current and future movements can be independently represented in the motor system. In monkeys trained to perform compound two arm reaching sequences, motor cortex activity could be decomposed into activity related to the ongoing and future reach (Zimnik and Churchland, 2021). Thus, the rapid execution of compound reaches did not depend on planning for the two reaches as a whole, rather on the ability to plan for the next reach while the current was under the way. Our model extends this idea of temporally overlapping, yet independent, planning processes to multiple future movements.

Of course, the presence of co-articulation implies that these different planning processes can also influence each other. Further research is needed to clarify the extent and conditions under which planning processes in sequential movements are dependent and how they influence each other.

### Mechanism of memory activation and deactivation

As previously discussed, it is crucial for the brain to flexibly generate sequences of movements in response to changes in environment (Rosenbaum, 2009). This flexibility requires a mechanism for activating sequence memories when appropriate and deactivating them when they no longer apply. Wolpert and Kawato (Wolpert and Kawato, 1998) introduced a responsibility estimator, a mechanism that assigns responsibility to different motor modules and dynamically adjusts their influence based on the accuracy of their output. Similar frameworks for responsibility assignment have been proposed in many other studies (Jacobs *et al*., 1991; Jordan and Jacobs, 2005; Heald, Lengyel and Wolpert, 2021).

In our task, we investigated how quickly memory traces can become deactivated or reactivated in response to a violation within a learned sequence. One behavioral observation was the absence of slowdown in performance prior to the violated press. This suggested that the memory trace for the sequence was not deactivated in anticipation of the violation, but only once it was encountered. A similar dependence of a memory on past, but not future, has been reported for partial repetition of finger press sequences (Shahbazi, Pruszynski and Diedrichsen, 2025) and arm reach sequences (Kashefi, Diedrichsen and Pruszynski, 2025).

Conversely, after the violated press, the performance benefit of the memory trace did not immediately resume. Instead, it took 3-4 correct presses before the memory trace was reactivated. This finding aligns with the notion that several movements compatible with a sequence memory must be experienced before that sequence memory can be (re-)activated (Kashefi, Diedrichsen and Pruszynski, 2025; Shahbazi, Pruszynski and Diedrichsen, 2025). Together, these findings point to a key constraint in sequential memory traces: activation or deactivation depends on the immediate history of compatible past elements, while future compatibility—despite evidence that future movements are planned concurrently with current movements—does not seem to trigger activation or deactivation.

### Neural control of sequential movements

Despite significant advances in understanding the neural architectures underlying skilled movements, a comprehensive neural framework explaining the computations during sequential movements is lacking. Our work addresses this gap by proposing a computational framework in which sequential finger presses are represented as a series of decision-making processes.

Previous studies have established links between neural activity in perceptual decision-making tasks and drift-diffusion models (Hanes and Schall, 1996; Gold and Shadlen, 2000; Roitman and Shadlen, 2002; Steinemann *et al*., 2024). More recently (Genkin *et al*., 2025), characterized how these decision-making computations manifest in the population dynamics of the premotor cortex. Building on this foundation, we extend the concept to sequential decision-making in motor behavior.

We propose that during the parallel preparation of multiple presses (e.g. the next three presses in a sequence), each upcoming press is represented by its own decision process unfolding within an orthogonal subspace of the neural population activity. This orthogonality aligns with our behavioral evidence for the independent planning of upcoming presses. Importantly, this orthogonal subspace framework raises a key question: as the current press is selected and executed, how does neural activity shift or rotate within this high-dimensional space, such that the subspace representing future 2 (two presses ahead) transitions into the future 1 subspace, and future 1 transitions into the subspace for the immediate next press? Further studies are needed to uncover the mechanisms underlying this dynamic transformation.

## Methods

### Participants

Twenty-six participants were recruited for the three experiments. Experiment 1 had fourteen participants (7 female, 7 male; mean age 26.3, SD 4.1), Experiment 2 twelve participants (7 female, 5 male; mean age 26.3, SD 5.6), and Experiment 3 twelve participants (8 female, 4 male; mean age 23.7, SD 3.4). For each participant, handedness was evaluated using the Edinburgh Handedness Inventory (Oldfield, 1971). According to this measure, all participants were right-handed, with an average handedness score of 82 ± 3. None of the participants reported any neurological disorders. Participants were compensated at a rate of $15 per hour. All procedures for this study were approved by the Health Science Research Ethics Board at the University of Western Ontario.

### Apparatus

A custom-made right-handed keyboard with force transducers under each of the five keys (Honeywell FS Series) was used. The transducers measured the isometric force produced by each finger, recorded at a rate of 200 Hz. A press was detected if the force exceeded the force threshold of 1.5 N, and a release was detected when the force went below 0.8 N.

Participants were seated in front of a computer screen at ~xxx cm distance, which displayed the static, white outline of a rectangle (stimulus rectangle) on the upper half of the screen against a plain, black background. Numerical stimuli making up the sequences were horizontally aligned (left to right) inside the stimulus rectangle, displaying digits from 1-5. Individual digits were 1.5 cm wide. Forces for each of the five fingers were displayed in the lower half of the screen as five lines that moved dynamically with the isometric force detected by the transducers. A red rectangle at the bottom of the display indicated an upper force limit of 0.5N and instructed participants to avoid exceeding this limit before the onset of a trial.

### General Procedures

In all experiments, participants performed sequences of finger presses in response to numerical cues displayed on a screen. Each finger of the right hand corresponded to a specific cue: thumb (1), index (2), middle (3), ring (4), and pinky (5). Participants were instructed to perform sequences as quickly and as accurately as possible while maintaining a minimum accuracy of 80%. Accuracy was measured as the percentage of sequences executed without error.

At the beginning of each trial, participants positioned their right hand on a custom keyboard-like device. Between trials, they were required to keep their finger forces within a red rectangle displayed on the screen. After a variable delay depending on the experiment, numerical cues appeared on the screen. A limited number of cues were initially visible from the left (referred to as the *visible horizon*), while the remaining were masked by asterisks. Each subsequent cue was revealed immediately following a press. A press was registered once the force applied to a finger exceeded a defined threshold. Immediate visual feedback followed each press: correct presses turned the corresponding cue green, and incorrect press turned it red **(Fig. 1B)**. Trials had to be completed within 10 seconds. Incorrect trials were signaled by a low-pitch auditory tone. The next trial started after a fixed 3000 ms inter-trial interval (ITI).

To motivate participants, they were awarded points for each trial based on their speed and accuracy: 0 points for trials with one or more press errors, +1 point for error-free trials, +2 points if execution time (ET) was within 105% of a personalized time threshold, +3 points if ET was within 98% of the threshold, and −1 point for failing to complete a trial within 10 seconds. Points were displayed immediately after the trial. The time threshold was designed to get increasingly difficult adjusting to each participant’s performance and was updated to the median ET of a block if it was faster than the participant’s best previous median (and block accuracy exceeded 80%). At the end of each block, participants were shown their total points, median ET, and error rate.

### Experiment 1 Procedures

This experiment was designed to measure integration of memory and sensory information in sequence production. Participants performed 7-digit sequences chosen to contain no runs of three (e.g. 123) or repetitions of the same digit (e.g. 22). Each sequence included all five fingers at least once. For each participant, two unique sequences were assigned as *repeating* sequences and appeared more frequently throughout the experiment. These sequences were identified by the symbols $ and #, while randomly generated sequences were identified with %.

At the beginning of each trial, during a random 600-1000 ms preparation time, a sequence type symbol was shown without numerical stimuli. The symbol was centered directly above the stimulus rectangle on the screen. Then the numerical cues appeared accompanied by a high-pitch tone that signaled participants to perform the sequence.

Each trial was chosen to have a *visible horizon* of either 1, 2, 3, or 7 (**Fig. 1B**). In *masked* trials, numerical cues remained covered by asterisks throughout the trial. Feedback was still provided with asterisks turning green or red based on the correctness of the press (**Fig. 1B**). Masked trials were used only for repeating sequences and forced participants to perform the sequence from memory.

The experiment consisted of four sessions conducted on separate days. The first three sessions served as training. In each training session, participants performed twelve blocks of 24 trials. Within each block, one-third of trials consisted of random sequences (%), one-third the first repeating sequence (#), and one-third the second repeating sequence ($). For the repeating sequences, half of the trials were masked (n = 4), and half unmasked. In unmasked trials (both repeating and random), visible horizons were counter-balanced (**Fig. 1C**). Trial order within each block was randomized.

In the fourth session, *violation* trials were introduced. In these trials, one of the repeating sequence symbols (# or $) was shown, but a single numerical cue at position 3, 4, or 5 was altered (**Fig. 2C**). Participants performed thirteen blocks of 24 trials. The first block was a warm-up block, identical in structure to those in the training sessions. Violation trials were then introduced in the remaining twelve blocks. The twelve blocks were grouped into two sets of six blocks (blocks 2-7 and 8-13). Each set contained 120 standard trials (with the same structure as in training) and 24 occasional violation trials.

### Experiment 2 Procedures

This experiment was designed to investigate in more detail how participants responded to conflicts between memory and sensory information (i.e., violations). Participants were divided into two groups; each assigned two repeating sequences that appeared more frequently throughout the experiment. All sequences were 11 digits long chosen to contain no runs of three (e.g. 123) or repetitions of a digit (e.g. 22).

At the beginning of each trial, a white fixation cross was placed inside the stimulus rectangle, positioned just left to the upcoming numerical cues. The numerical cues then appeared with a chosen visible horizon of 2, 3, 4, or 11. During an interval of 1500 ms, participants were instructed to maintain fixation on the cross. The onset of execution was signaled by the cross turning green, accompanied by a high-pitched tone.

The experiment was conducted across three sessions. The first session served as training, during which participants practiced their assigned repeating sequences across six blocks of 32 trials. The subsequent two sessions served as test and included random, repeating, and violation trials. In violation trials a repeating sequence with a single altered cue at positions 5, 7, or 9 was used. Each test session consisted of eight blocks of thirty trials. Within every set of four blocks, 40% were random, 40% repeating, and 20% violation trials.

### Experiment 3 Procedures

This experiment was designed to investigate how sequence memory was deactivated when a sequence became inconsistent with a learned sequence. As in Experiment 2, participants were divided into two groups; each assigned two repeating sequences of length 11. Like in Experiment 2, participants were instructed to fixate on the fixation cross, after which numerical cues appeared to the right of it. However, unlike in Experiment 2, participants could start execution as soon as numerical cues appeared. This was accompanied by the cross turning green and a high-pitched sound.

The experiment consisted of three sessions with the same structure as Experiment 2. The only difference was that in violation trials, all cues following a specific position (either digit 5, 7, or 9) were altered from the repeating sequence.

### Data Analysis

Participants’ performance was measured using movement times (MTs), inter-press intervals (IPIs), and error rates (ERs). *Movement time* was defined as the time between the first press and the release of the last press in the sequence (i.e. the time between the first and the last crossing of the force threshold; **Fig. 1A**). inter-press intervals (IPIs) were defined as the time between successive presses in the sequence (**Fig. 1A**). Error rates (ERs) were defined as the proportion of incorrect trials (i.e. trials with one or more incorrect presses) within each block. For MT and IPI analysis, subject-specific values were computed as the median measure of all trials within each condition. Trials containing errors were excluded from MT and IPI analysis.

### Decision Model

For a single press, we modeled the decision of which finger to press as evidence accumulation to a threshold T (Usher and McClelland, 2001; Bogacz *et al*., 2006). The evidence at time *t* was represented by a 5-dimensional vector *X*_*t*_, with each component corresponding to one finger.

The accumulation process was driven by two sources of input: sensory input *S* from the visible numerical cue, and memory input *M* learned from prior training. Both *S* and *M* were one-hot encoded vectors indicating the relevant finger. These inputs were integrated into a total input vector:

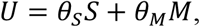

where *θ*_*S*_ and *θ*_*M*_were the respective input weights for sensory and memory. At each time step (Δt = 1 ms), the evidence vector was updated according to:

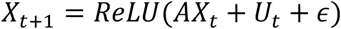

where ∈ ~ *N*(0, σ^2^*I*) represented Gaussian noise independently and identically applied to each finger. *A* is a 5 × 5 state transition matrix encoding self and lateral inhibition. This was realized by setting its diagonals to 1 − *α* for self-inhibition, and off-diagonals to −*β* for lateral inhibition where *α* and *β* were bounded between 0 and 1. The ReLU function ensured non-negative evidence values: *ReLU*(*x*) = max (0, *x*).

### Fitting Single-Press Parameters

To fit the model to the data, we took a stage-wise approach. In the first stage, we fit the model parameters for a single press using the Horizon 1 trials from Experiment 1. In this condition, the memory input for the current press started right after the previous press, and the sensory input after a delay of 90 ms. Model fitting was performed in three sequential steps: First, we fitted the threshold (*T*) by minimizing the squared error between model-predicted and observed average reaction times in random trials (*M* = 0). Second, we fitted the memory weight (*θ*_M_) by minimizing the squared error for repeating trials (*M* = *S*). Finally, we fitted the lateral inhibition constant (*β*) by minimizing the squared error for violation trials (*M* ≠ *S*). The parameters *θ*_S_, σ, and *α* were predefined to reduce the parameter space. Fitting was performed using the swarm optimization algorithm (Kennedy and Eberhart, 2002). Only presses at violation positions (2, 3, or 4) were used for fitting.

### Modeling Sequences

To model sequences, we extended the single-press model by assuming that up to three future presses were planned *in parallel* based on previous findings (Ariani *et al*., 2021). Here, planning is defined broadly as encompassing all processes from the initial stimulus processing to the execution of a press, collectively reflected in the reaction time to a cue. Each press’s accumulation process operated *independently* with decreasing input weights for future presses:

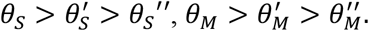

For masked digits, the sensory input was set to zero.

### Fitting Sequence Parameters

In the second stage of model fitting, we used the data from the Horizon > 1 conditions in Experiment 1. Parameters estimated in the first stage (σ, *T, α, β, θ*_S_, *θ*_M_) were held fixed, and we optimized only the input weights for future presses. Fitting was performed in two steps: First, we estimated the future sensory weights (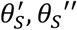) by maximizing the Pearson correlation between model-predicted and observed average movement times (MT) in random trials across horizons. Next, we estimated the future memory weights (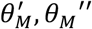) by maximizing the same metric in repeating trials. Parameters were fit using swarm optimization. The fitted model was then used to predict performance in violation trials with Horizon > 1.

### Modeling Memory Deactivation and Reactivation

To model the behavior on presses following the violation press, we hypothesized that memory input weights (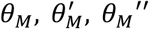) were dynamically updated in response to the violation. Memory deactivation (*τ*_*deact*_) and memory reactivation (*τ*_*react*_) were represented by two time-constants with values between zero and one. When a conflict occurred (*M* ≠ *S*), memory input weights were reduced according to:

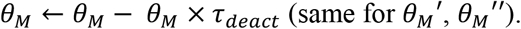

When the conflict resolved (*M* = *S*), memory weights recovered toward their original value:

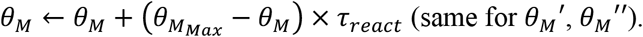

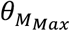 indicated the original memory weight before the conflict.

### Fitting Memory Deactivation and Reactivation Parameters

To estimate *τ*_*deact*_ and *τ*_*react*_, we used the violation trials from Experiment 2 in Horizon > 1 conditions. For each inter-press interval, we calculated the relative difference between average values in repeating and violation trials, which served as the target behavior for model fitting. *τ*_*deact*_ and *τ*_*react*_ were optimized to minimize the sum of squared errors between predicted and observed relative differences across all inter-press intervals. The optimization was performed individually per participant using swarm optimization. The resulting estimates were averaged to obtain population level parameters. The population *τ*_*deact*_ parameter was used to predict the performance on violation trials in Experiment 3.

### Statistical Analysis

We used within-subject design across all experiments. Statistical analyses were conducted using Statsmodels 0.14.4 (Seabold and Perktold, 2010). For each test, we report degrees of freedom, test statistics, and p-values. Two-way and one-way repeated measures analysis of variance (ANOVA) and paired t-tests were employed. In Experiment 1, the factors were visible horizon (five levels: masked, H1, H2, H3, H7), sequence type (three levels: repeating, violation, random), and session (three levels: Day 1, Day 2, Day 3). In Experiment 2 and 3, the factors were visible horizon (four levels: H2, H3, H4, H11), and sequence type (three levels: repeating, violation, random). All t-tests were two-sided.

## Acknowledgements

This work was supported by a project grant from the Canadian Institutes of Health Research (CIHR, PJT-175010) to A.P. and J.D. and the Canada First Research Excellence Fund (BrainsCAN) to Western University. AP received a salary award from the Canada Research Chairs program.

## Competing Interest

The authors declare no competing interests.

## Supplementary Figures

**Supplementary Figure 1.**
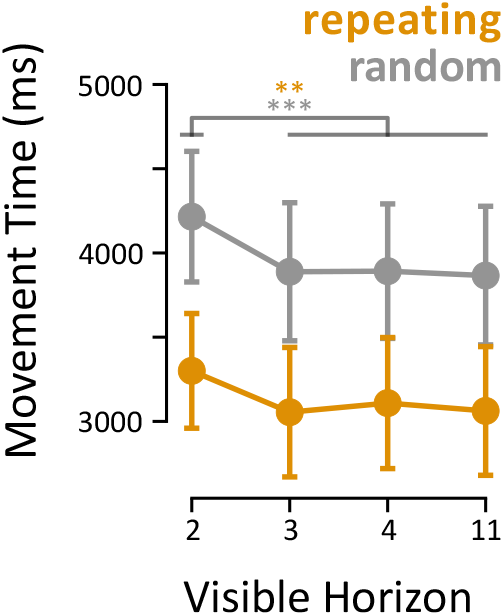
Group-averaged movement times for each visible horizon in repeating (yellow) and random (gray) conditions in Experiment 2. Error bars show the standard error of the mean. ** and *** show *p* < 0.01 and *p* < 0.001.

## Notes

### Competing Interest Statement

The authors have declared no competing interest.

## References

Abrahamse, E.L. et al. (2013) “Control of automated behavior: insights from the discrete sequence production task,” Frontiers in human neuroscience, 7, p. 82.

Adams, J.A. (1987) “Historical review and appraisal of research on the learning, retention, and transfer of human motor skills,” Psychological bulletin, 101(1), pp. 41–74.

Ariani, G. et al. (2021) “The Planning Horizon for Movement Sequences,” eNeuro, 8(2). Available at: 10.1523/ENEURO.0085-21.2021.

Bashford, L. et al. (2022) “Motor skill learning decreases movement variability and increases planning horizon,” Journal of neurophysiology, 127(4), pp. 995–1006.

Berniker, M. and Kording, K. (2011) “Bayesian approaches to sensory integration for motor control,” Wiley interdisciplinary reviews. Cognitive science, 2(4), pp. 419–428.

Bogacz, R. et al. (2006) “The physics of optimal decision making: a formal analysis of models of performance in two-alternative forced-choice tasks,” Psychological review, 113(4), pp. 700–765.

Botvinick, M. et al. (1999) “Conflict monitoring versus selection-for-action in anterior cingulate cortex,” Nature, 402(6758), pp. 179–181.

Fitts, P.M. (1964) “Perceptual-Motor Skill Learning,” in Categories of Human Learning. Elsevier, pp. 243–285.

Fitts, P.M. and Posner, M.I. (1979) Human Performance. Westport, CT: Praeger.

Genkin, M. et al. (2025) “The dynamics and geometry of choice in the premotor cortex,” Nature, pp. 1–9.

Gold, J.I. and Shadlen, M.N. (2000) “Representation of a perceptual decision in developing oculomotor commands,” Nature, 404(6776), pp. 390–394.

Hanes, D.P. and Schall, J.D. (1996) “Neural control of voluntary movement initiation,” Science (New York, N.Y.), 274(5286), pp. 427–430.

Heald, J.B., Lengyel, M. and Wolpert, D.M. (2021) “Contextual inference underlies the learning of sensorimotor repertoires,” Nature, 600(7889), pp. 489–493.

Hikosaka, O. et al. (1999) “Parallel neural networks for learning sequential procedures,” Trends in neurosciences, 22(10), pp. 464–471.

Howard, J.H. and Howard, D.V. (1997) “Age differences in implicit learning of higher order dependencies in serial patterns,” Psychology and aging, 12(4), pp. 634–656.

Jacobs, R.A. et al. (1991) “Adaptive mixtures of local experts,” Neural computation, 3(1), pp. 79–87.

Jiménez, L. and Méndez, C. (1999) “Which attention is needed for implicit sequence learning?,” Journal of experimental psychology. Learning, memory, and cognition, 25(1), pp. 236–259.

Jordan, M.I. and Jacobs, R.A. (2005) “Hierarchical mixtures of experts and the EM algorithm,” in Proceedings of 1993 International Conference on Neural Networks (IJCNN-93-Nagoya, Japan). 1993 International Conference on Neural Networks (IJCNN-93-Nagoya, Japan), IEEE, pp. 1339–1344 vol. 2.

Karni, A. et al. (1995) “Functional MRI evidence for adult motor cortex plasticity during motor skill learning,” Nature, 377(6545), pp. 155–158.

Kashefi, M. et al. (2024) “Future movement plans interact in sequential arm movements,” eLife, 13. Available at: 10.7554/elife.94485.3.

Kashefi, M., Diedrichsen, J. and Pruszynski, J.A. (2025) “Motor sequence learning involves better prediction of the next action and optimization of movement trajectories,” The Journal of neuroscience: the official journal of the Society for Neuroscience [Preprint]. Available at: 10.1523/JNEUROSCI.0299-25.2025.

Keele, S.W. et al. (1995) “On the modularity of sequence representation,” Journal of motor behavior, 27(1), pp. 17–30.

Kennedy, J. and Eberhart, R. (2002) “Particle swarm optimization,” in Proceedings of ICNN’95 - International Conference on Neural Networks. ICNN’95 - International Conference on Neural Networks, IEEE, pp. 1942–1948 vol.4.

Kent, R.D. and Minifie, F.D. (1977) “Coarticulation in recent speech production models,” Journal of phonetics, 5(2), pp. 115–133.

Krakauer, J.W. et al. (2019) “Motor Learning,” Comprehensive Physiology, 9(2), pp. 613–663.

Lewicki, P., Hill, T. and Bizot, E. (1988) “Acquisition of procedural knowledge about a pattern of stimuli that cannot be articulated,” Cognitive psychology, 20(1), pp. 24–37.

Logan, G.D. (1988) “Toward an instance theory of automatization,” Psychological review, 95(4), pp. 492–527.

Mizes, K.G.C. et al. (2023) “Dissociating the contributions of sensorimotor striatum to automatic and visually guided motor sequences,” Nature neuroscience, 26(10), pp. 1791–1804.

Oldfield, R.C. (1971) “The assessment and analysis of handedness: the Edinburgh inventory,” Neuropsychologia, 9(1), pp. 97–113.

Picard, N., Matsuzaka, Y. and Strick, P.L. (2013) “Extended practice of a motor skill is associated with reduced metabolic activity in M1,” Nature neuroscience, 16(9), pp. 1340–1347.

Proctor, R.W. and Dutta, A. (1995) Skill Acquisition and Human Performance. Thousand Oaks, CA: SAGE Publications (Advanced Psychology Text Series).

Ramkumar, P. et al. (2016) “Chunking as the result of an efficiency computation trade-off,” Nature communications, 7, p. 12176.

Roitman, J.D. and Shadlen, M.N. (2002) “Response of neurons in the lateral intraparietal area during a combined visual discrimination reaction time task,” The Journal of neuroscience: the official journal of the Society for Neuroscience, 22(21), pp. 9475–9489.

Rosenbaum, D.A. (2009) Human Motor Control. 2nd ed. San Diego, CA: Academic Press.

Rosenbaum, D.A., Kenny, S.B. and Derr, M.A. (1983) “Hierarchical control of rapid movement sequences,” Journal of experimental psychology. Human perception and performance, 9(1), pp. 86–102.

Schvaneveldt, R.W. and Gomez, R.L. (1998) “Attention and probabilistic sequence learning,” Psychological research, 61(3), pp. 175–190.

Seabold, S. and Perktold, J. (2010) “Statsmodels: Econometric and statistical modeling with python,” in Proceedings of the Python in Science Conference. Python in Science Conference, SciPy, pp. 92–96.

Shahbazi, M., Pruszynski, J.A. and Diedrichsen, J. (2025) “Repetition effects reveal the subsequence representation of actions,” Journal of neurophysiology, 134(2), pp. 691–697.

Simó, L.S., Krisky, C.M. and Sweeney, J.A. (2005) “Functional neuroanatomy of anticipatory behavior: dissociation between sensory-driven and memory-driven systems,” Cerebral cortex (New York, N.Y.: 1991), 15(12), pp. 1982–1991.

Steinemann, N. et al. (2024) “Direct observation of the neural computations underlying a single decision,” eLife, 12. Available at: 10.7554/eLife.90859.

Trommershäuser, J., Maloney, L.T. and Landy, M.S. (2008) “Decision making, movement planning and statistical decision theory,” Trends in cognitive sciences, 12(8), pp. 291–297.

Usher, M. and McClelland, J.L. (2001) “The time course of perceptual choice: the leaky, competing accumulator model,” Psychological review, 108(3), pp. 550–592.

Vassena, E., Deraeve, J. and Alexander, W.H. (2020) “Surprise, value and control in anterior cingulate cortex during speeded decision-making,” Nature human behaviour, 4(4), pp. 412–422.

Verwey, W.B. (1996) “Buffer loading and chunking in sequential keypressing,” Journal of experimental psychology. Human perception and performance, 22(3), pp. 544–562.

Verwey, W.B. (2001) “Concatenating familiar movement sequences: the versatile cognitive processor,” Acta psychologica, 106(1–2), pp. 69–95.

Welford, A.T. (1968) Fundamentals of skill. London, England: Methuen young books (Manual of Modern Psychology S.).

Willingham, D.B. (1998) “A neuropsychological theory of motor skill learning,” Psychological review, 105(3), pp. 558–584.

Willingham, D.B. et al. (2000) “Implicit motor sequence learning is represented in response locations,” Memory & cognition, 28(3), pp. 366–375.

Wolpert, D.M. and Kawato, M. (1998) “Multiple paired forward and inverse models for motor control,” Neural networks: the official journal of the International Neural Network Society, 11(7–8), pp. 1317–1329.

Wolpert, D.M. and Landy, M.S. (2012) “Motor control is decision-making,” Current opinion in neurobiology, 22(6), pp. 996–1003.

Wong, A.L. et al. (2015) “Explicit knowledge enhances motor vigor and performance: motivation versus practice in sequence tasks,” Journal of neurophysiology, 114(1), pp. 219–232.

Wu, T., Kansaku, K. and Hallett, M. (2004) “How self-initiated memorized movements become automatic: a functional MRI study,” Journal of neurophysiology, 91(4), pp. 1690–1698.

Wymbs, N.F. and Grafton, S.T. (2015) “The human motor system supports sequence-specific representations over multiple training-dependent timescales,” Cerebral cortex (New York, N.Y.: 1991), 25(11), pp. 4213–4225.

Xu, D. et al. (2022) “Cortical processing of flexible and context-dependent sensorimotor sequences,” Nature, 603(7901), pp. 464–469.

Zimnik, A.J. and Churchland, M.M. (2021) “Independent generation of sequence elements by motor cortex,” Nature neuroscience, 24(3), pp. 412–424.

